# Lipid specificity of action of SARS-CoV-2 fusion peptide fragments on model membranes

**DOI:** 10.1101/2024.12.11.627892

**Authors:** Egor V. Shekunov, Pavel E. Volynsky, Svetlana S. Efimova, Elena T. Aliper, Roman G. Efremov, Olga S. Ostroumova

**Affiliations:** Institute of Cytology of Russian Academy of Sciences, Tikhoretsky 4, 194064, Saint Petersburg, Russian Federation; Shemyakin-Ovchinnikov Institute of Bioorganic Chemistry of Russian Academy of Sciences, Miklukho-Maklaya Str. 16/10, 117997, Moscow, Russian Federation

**Keywords:** COVID-19, SARS-CoV-2, Lipid Bilayers, Fusion Peptide, Membrane Fusion

## Abstract

The study focuses on investigating the interaction of SARS-CoV-2 fusion peptide fragment with model membranes of various lipid composition to elucidate the molecular mechanisms of peptide-derived membrane fusion. The work utilized the short fragment of SARS-CoV-2 fusion peptide which is homologous to 816-827 region of the native SARS-CoV-2 FP (FP_816-827_) and contains the highly conserved LLF motif responsible for membrane fusion, and its ineffective analogue (mFP_816-827_), where LLF motif was replaced for AAA. Using fluorescence fusion assay, it was demonstrated that the LLF motif plays a key role in inducing liposome fusion, whereas its replacement completely abolishes this capability. The fusogenic activity of the peptide strictly depended on the vesicle lipid composition. It was potentiated by phosphatidylethanolamine and inhibited by phosphatidylserine. Molecular dynamics revealed that both peptides predominantly adopt an α-helical conformation; however, the native peptide interacts more strongly with the hydrophobic core of the membrane by increasing peptide-lipid hydrophobic contacts, while the mutant version exhibits a more superficial localization. Differential scanning microcalorimetry data indicated that the ability of FP_816-827_ to disturb lipid packing increased with decreasing membrane lipid tail length. The molecular mechanisms underlying the fusogenic activity of the SARS-CoV-2 fusion peptide were identified, specifically its ability to cluster phospholipid head groups in its own vicinity. As a result, local regions with positive spontaneous curvature are formed in the outer monolayer, facilitating membrane fusion. The findings highlight the role of membrane composition and lipid architecture in the mechanism of viral fusion with host cells.

## Introduction

SARS-CoV-2, also known as Severe Acute Respiratory Syndrome Coronavirus 2, is a novel coronavirus accountable for the global COVID-19 pandemic. It belongs to the *Coronaviridae* family and shares close genetic kinship with other coronaviruses, including SARS-CoV and MERS-CoV [1]. The cumulative count of COVID-19 cases reported on 31 March 2024 is equal to 775,251,779 [2]. While a considerable proportion of cases manifest with mild respiratory symptoms or remain asymptomatic, severe disease manifestations such as Acute Respiratory Distress Syndrome (ARDS), multiorgan failure, and fatalities can occur. Particularly vulnerable populations include the elderly patients and individuals with compromised immune function [3].

SARS-CoV-2 represents a single-stranded, positive-sense RNA virus enveloped within a lipid bilayer, adopting a spherical configuration. Analogous to other enveloped viruses, the life cycle of SARS-CoV-2 initiates with its penetration into the target cell. To accomplish this, the virus must instigate the fusion of its lipid envelope with the cellular membrane, thereby facilitating the ingress of its genetic material into the cell’s cytoplasm. The recognition of virus cellular receptors and the fusion with the host cell membrane are implemented by the spike protein (S) [4]. This glycoprotein belongs to the class I fusion protein and consists of 1273 amino acid sequence, divided into two subunits, namely S1 (14-685) and S2 (686-1273) [5]. For the proper execution of its functions by the S2 subunit, the original S protein necessitates cleavage by cellular proteases. Two distinct cleavage sites (S1/S2 and S2’) can be activated by a broad spectrum of proteases [6].

The S1 subunit consists of the N-terminal domain (NTD) and the receptor-binding domain (RBD) [7]. The primary function of the S1 subunit resides in engagement with target cells via interactions with specific receptor, human angiotensin-converting enzyme 2 (ACE2) receptor. The S2 subunit comprising the fusion peptide (FP), two heptad repeats (HR1 and HR2), a transmembrane domain (TMD), and a cytoplasmic domain (CPD), implements the fusion event between the viral particle and the host cell [5]. Fusion initiation involves the activation of the FP domain, which is inserted into the target membrane. This interaction initiates cellular-viral contact, culminating in the formation of a six-helix bundle (6-HB) structure, constituted by a trimeric complex comprising HR1 and HR2 antiparallel assemblies [4].

It is recognized that the fusion peptide (FP) structurally comprises two subdomains, FP1 and FP2, yet the precise localization of this domain and the mechanism of peptide-induced fusion remain subjects of ongoing discussion. Several conserved sequences within *Coronaviridae* have been proposed as potential candidates for fulfilling the FP domain role in the S protein of SARS-CoV-2, namely 770-788 and 864-886 [8,9]. However, the prevailing consensus in the majority of studies suggests that the 816-833 region corresponds to the FP1 region [10], while 834-855 fragment constitutes the FP2 subdomain [6]. FP1 exhibits a higher alpha-helicity compared to FP2 and harbors a highly hydrophobic LLF motif, a feature conserved across the *Coronaviridae* family [11]. FP1 is presumed to insert into the target membrane, perturbing its architecture and thereby instigating fusion [12]. However, the precise functional role of FP2 remains incompletely elucidated. Notably, investigations with SARS-CoV have revealed that FP2 induces enhanced membrane ordering and interacts with FP1, forming an elongated fusion peptide FP1-FP2 with augmented ordering activity. The degree of membrane ordering reflects the dehydration effect, which plays a crucial role in fusion. Consequently, it is speculated that this may describe the degree of fusogenic activity of peptides [6]. Similar observations demonstrating a heightened fusion-inducing capacity of FP1-FP2 have been described for SARS-CoV-2 [39]. Niort et al. has demonstrated that SARS–CoV-2 FP2 is unable to induce fusion, and FP1–FP2 is not characterized by increased fusogenic activity [13].

The efficacy of fusion mediated by viral proteins is also contingent upon the lipid composition of the merging membranes, as highlighted in several studies [14–16]. Phosphatidylethanolamine (PE), characterized by its inverted cone geometry [17,18], is believed to have a pivotal role in facilitating fusion by diminishing the energy barrier required for initiating an fusion intermediate called stalk [19–21]. Charged lipids also play an important role in fusion. For example, it has been suggested that phosphatidylserine (PS) in the membranes of target cells promotes the attraction of fusion peptides having a positive charge [22,23]. Additionally, the integrity of lipid rafts has been shown to modulate the activity of fusion peptides of HIV-1 [24], Ebola [25], influenza A [26], and SARS-CoV-2 [27,28]. Furthermore, the addition of pharmacological agents targeting cholesterol, such as cholesterol 25-hydroxylase, has been demonstrated to suppress the development of SARS-CoV-2 [29]. The existence of conflicting data regarding the accurate amino acid sequence of the fusion peptide in SARS-CoV-2, incomplete insights into the architectural rearrangements of membrane structures upon interaction with viral fusion proteins, and a limited understanding of the molecular mechanisms underpinning membrane fusion indicate the need for further investigation. Furthermore, studying the specific interactions occurring in the membrane between viral fusion peptides and lipids can help in the development of innovative antiviral strategies and drugs [13,30,31].

Studies employing synthetic peptides related to distinct viral FP domains and model lipid membranes provide valuable insights, enabling a precise focus on molecular features of lipid matrix rearrangement during fusion [32,33]. Previous studies of the fusogenic activity of the SARS-CoV-2 fusion peptide demonstrated the critical importance of the LLF domain for membrane fusion [6,34]. Substituting this region with three consecutive alanine residues results in the loss of the protein’s ability to induce fusion [35]. In this study, we explored the fusogenic activity of synthetic peptides homologous to the region 816-827 (_816_SFIEDLLFNKVT_827_; FP1) and its analogue with a mutation in the highly conserved motif LLF (_816_SFIEDAAANKVT_827_) of the SARS-CoV-2 S protein and how the replacement of lipid head groups, as well as the length of their hydrophobic segments, affects the function of the peptide. Furthermore, we examined the molecular mechanisms underlying the functional activity of the SARS-CoV-2 fusion peptide. To achieve this, we employed methods such as membrane fusion assays, differential scanning microcalorimetry, and molecular dynamics simulations.

## Results and Discussion

### Fusogenic activity of SARS-CoV-2 FP1 and FP2 fragments in membranes of various composition

We investigated the ability of short fragments of SARS-CoV-2 FP1 (_816_SFIEDLLFNKVT_827_ (FP_816-827_)) to trigger the fusion of membranes of various lipid compositions. The peptide, _816_SFIEDAAANKVT_827_ (mFP_816-827_), which is homologous to 816-827 region of the native SARS-CoV-2 FP with the highly conserved LLF motif responsible for membrane fusion replaced for AAA, was used as a negative control [6,34,35]. The activity and functions of many membrane proteins can be regulated by the lipids constituting the membrane. This regulatory activity can be specific, arising from selective lipid-protein interactions, or nonspecific, driven by the physical properties of lipids, including the charge of the head group, the length and order of the hydrocarbon tail, the geometry of the lipid, and the lateral tension [36].

Figure 1 demonstrates the calcein leakage due to fusion of POPC/SM/CHOL (60/20/20 mol.%) (a), POPC/POPE/SM/CHOL (30/30/20/20 mol.%) (b) and POPS/SM/CHOL (60/20/20 mol.%) (c) liposomes induced by 150 µM of various tested fragments. A peptide homologous to the FP1 region (FP_816-827_) induced membrane fusion, and its efficiency strictly depended on the lipid composition of the vesicles. As expected, the addition of mFP_816-827_, in which the LLF motif responsible for the fusogenic activity of coronaviruses [6,35] was replaced by AAA, did not cause marker leakage in all tested lipid systems (*IF* did not exceed 10% independently on lipid composition) (Figure 1). This indicates the significant role of the LLF sequence in the fusogenic function implementation by the SARS-CoV-2 fusion peptide. The mean values of maximum marker leakage caused by peptide-induced fusion of lipid vesicles are summarized in Table 1.

**Figure 1.**
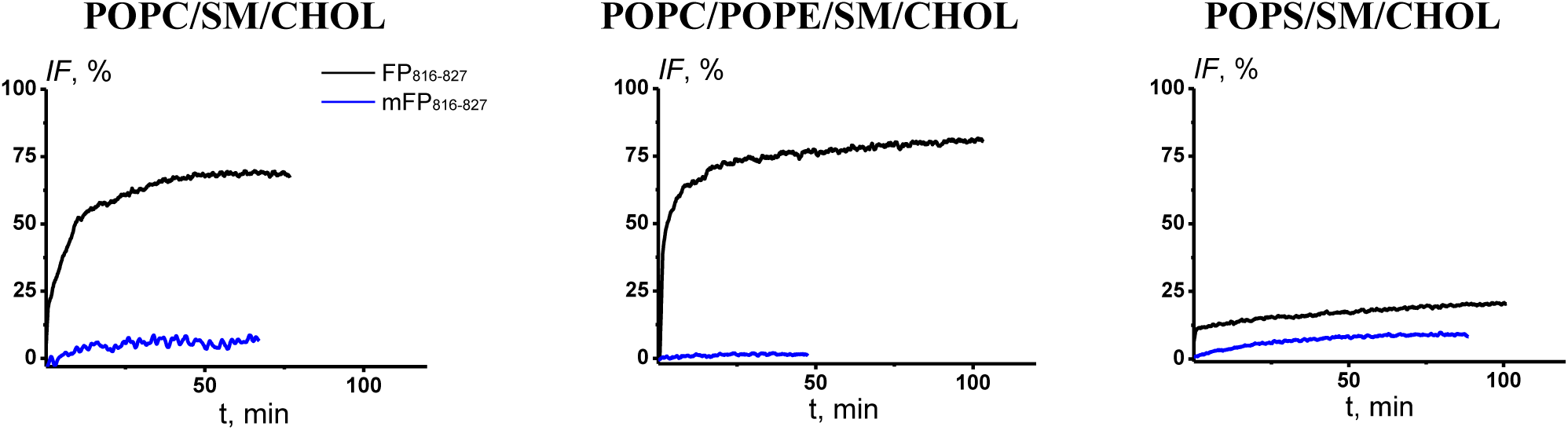
The time dependence of relative fluorescence of calcein leaked from POPC/SM/CHOL (60/20/20 mol.%), POPC/POPE/SM/CHOL (30/30/20/20 mol.%), and POPS/SM/CHOL (60/20/20 mol.%) vesicles in the presence of 150 μM FP_816-827_ and mFP_816-827_. The relationship between the color and type of peptide is shown on the figure.

**Table 1.** The ability of different fragments of SARS-CoV-2 FP1 and FP2 to induce calcein leakage associated with the fusion of POPC/SM/CHOL (60/20/20 mol.%), POPC/POPE/SM/CHOL (30/30/20/20 mol.%) and POPS/SM/CHOL (60/20/20 mol.%) vesicles at 150 μM.

| FP fragment | POPC/SM/CHOL | POPC/POPE/SM/CHOL | POPS/SM/CHOL |
| --- | --- | --- | --- |
| FP<sub>816-827</sub><br>(<sub>816</sub>SFIEDLLFNKVT<sub>827</sub>) | 46 ± 14 | 67 ± 21 | 16 ± 8 |
| mFP<sub>816-827</sub><br>(<sub>816</sub>SFIEDAAANKVT<sub>827</sub>) | 8 ± 6 | 6 ± 4 | 6 ± 2 |

The fusogenic ability of FP_816-827_ peptides depended on the membrane lipid composition (Table 1). *IF* produced by addition of FP_816-827_ to suspension of POPC/SM/CHOL liposomes was moderate (45 %). The inclusion of POPE, a nonlamellar lipid of a conical shape [17] with the cross-sectional area of the hydrocarbon chains’ region exceeding the cross-sectional area of polar head, led to slight increase in the efficiency of the peptide (*IF* reached for 65%) (Table 1). The role of phosphatidylethanolamine in fusion has been studied extensively [19–21]. The enhancing effect of this lipid might be due to its ability to reduce the energy barrier that must be overcome for the successful formation of the central fusion intermediate, the stalk, by forming the inverted hexagonal lipid phase. It has been established that in membranes composed of dioleoylphosphatidylcholine the energy required to induce the stalk is approximately 45 *kT*. On the contrary, in membranes made from dioleoylphosphatidylethanolamine, the stalk energy is strongly negative, around of −30 *kT* [37]. This implies that for this lipid, stalk formation is energetically favorable and, therefore, promotes fusion. Replacement of neutral POPC for negatively charged POPS led to significant changes in the fusogenic activity of FP816-827 (15%) (Table 1). Thus, the incorporation of negatively charged lipid potentiated and inhibited the fusogenic activity of the peptide. This result was partially expected, given that the presence of glutamic and aspartic acids in the peptide structure imparted a net negative charge to FP_816-827_, leading to electrostatic repulsion between the peptide molecules and the POPS head groups. Additionally, this should be due to the difference between electrostatic interactions of C-terminus of the peptide with the carboxyl group of serine, because FP_816-827_ is flanked by polar T827. It is known, that the charge of lipid headgroups influences the conformation, orientation and penetration of the peptide. Molecular simulation data showed that in the presence of POPS, the percentage of α-helicity of FP_816-827_ was higher than in the presence of POPC (Figure 3). Moreover, FP_816-827_ demonstrated more pronounced effects on thermotropic behavior of phosphatidylserines than phosphatidylcholines with the chains of same lengths (Table 3, Table 4). The importance of the tilt angle of the C-terminal end of the HIV FP (gp-41) for its fusogenic activity was described by Haque et al. [38]. The charge of the polar lipid heads is not the single factor that was able to alter the fusogenic activity of FP_816-827_. The importance of charge-charge interactions between FP and membrane for the peptide fusion activity is described in [39]. The authors have demonstrated that the FP1-FP2 of SARS-CoV-2 (_816_-SFIEDLLFNKVTLADAGFIKQYGDCLGDIAARDLICAQKF-_855_) possessing a negative charge at neutral pH, exhibits low fusogenic activity. However, under acidic conditions, when the peptide charge becomes positive, its ability to induce membrane fusion is significantly increased.

### Molecular dynamics simulation of interaction of FP_816-827_ and mFP_816-827_ with membranes of various composition

To deep insight into different effects of POPC, POPE and POPS on the fusogenic activity of FP_816-827_ the molecular dynamics calculations were performed. For comparison, the interaction of mFP_816-827_ with lipid bilayers of various composition was also investigated by *in silico* approach. Firstly, coarse-grained simulations were performed for both peptides in each of the bilayer compositions (POPC/SM/CHOL (60/20/20 mol.%), POPC/POPE/SM/CHOL (30/30/20/20 mol.%), POPS/SM/CHOL (60/20/20 mol.%)), amounting to a total of 12 trajectories for each peptide. In each replica the peptide was arbitrarily oriented and placed in the aqueous environment away from the bilayer, after which 1000-ns simulations were run. Both FP_816-827_ and mFP_816-827_ sooner or later ended up bound to the bilayer in 4 out of 4 replicas in systems with POPC/SM/CHOL and POPC/POPE/SM/CHOL bilayers and in 3 out of 4 replicas in the case of the negatively charged POPS/SM/CHOL bilayers. Binding modes of the peptides averaged over all the replicas in which binding took place for each bilayer composition are presented in Figure 2. As is evident from Figure 2, both peptides position themselves in a similar manner in relation to the bilayer in all cases, the N-terminus submerging to a greater extent than the C-terminus, the orientation thus being tilted. Even at the level of coarse-grained simulations it is clear that the 821-LLF-823 motif in the FP_816-827_ greatly contributes towards the interaction with the bilayer, including its deeper portion made up by the hydrophobic tails of the lipids. Meanwhile, the 821-AAA-823 motif in the mFP_816-827_ interacts with the same stratum of the bilayer far more feebly.

**Figure 2.**
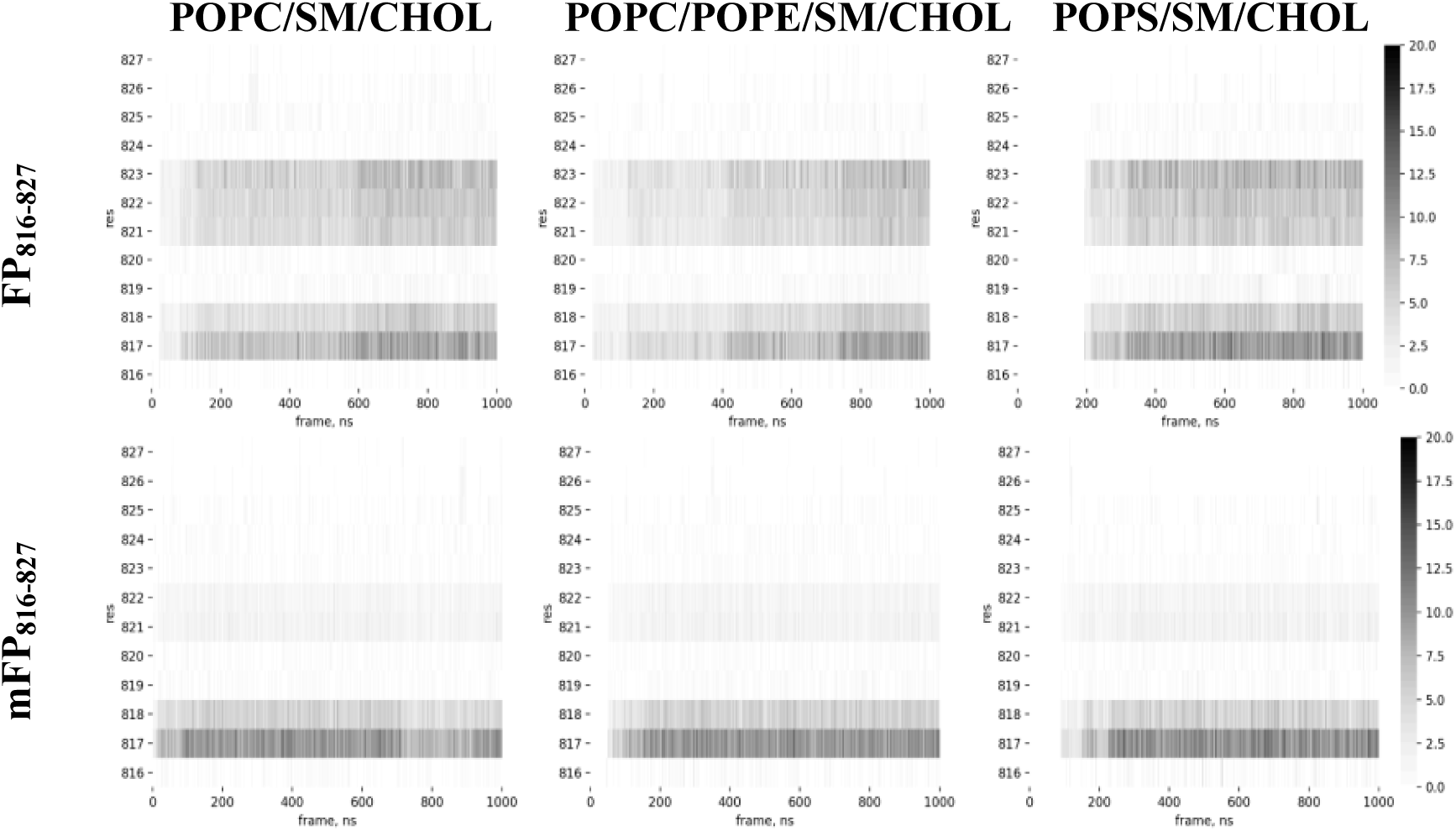
Binding modes of FP_816-827_ (above) and mFP_816-827_ (below) averaged over all the replicas for a given bilayer composition in which binding occurs (4 for POPC/SM/CHOL (60/20/20 mol.%) and POPC/POPE/SM/CHOL (30/30/20/20 mol.%), 3 for POPS/SM/CHOL (60/20/20 mol.%)) presented as heatmaps. Residue numbers are plotted against time frames of the trajectories, while the number of contacts made by the peptide with non-polar lipid tails is represented as a spectrum of shades of grey (from white as zero to black as 20, as indicated by the colour bar on the right).

Secondly, in the trajectories in which the peptide bound to the model bilayer, the final frames were converted from coarse-grained to all-atom model systems and additionally simulated for 500 ns each. In all cases, the peptides were somewhat mobile in relation to the starting state, as is evident from the RMSD values (Table S1, Supplementary materials), indicating that some mutual adjustment between the peptide and the environment formed by the membrane was taking place. In all cases, the N- and C-terminus showed some degree of mobility (data not shown) with respect to each other, indicating that the backbone of the peptide does undergo structural fluctuations across the trajectories. Indeed, the secondary structure of both peptides does not remain uniform throughout the simulations (Figure 3). For both FP_816-827_ and mFP_816-827_, the more C-terminal residues were less likely to remain in the α-helical conformation, but the rest of the peptide demonstrated considerable capacity for forming helices, sometimes even reverting to a helical state long after becoming unordered. Overall, one could characterize the helical state as labile and not fully resistant to destabilization. Both peptides were the most likely to remain in a helical state in POPS/SM/CHOL systems if one considers all helical conformations assumed (alpha-, 3- and 5-helix), most notably around the N-terminal and central portions.

**Figure 3.**
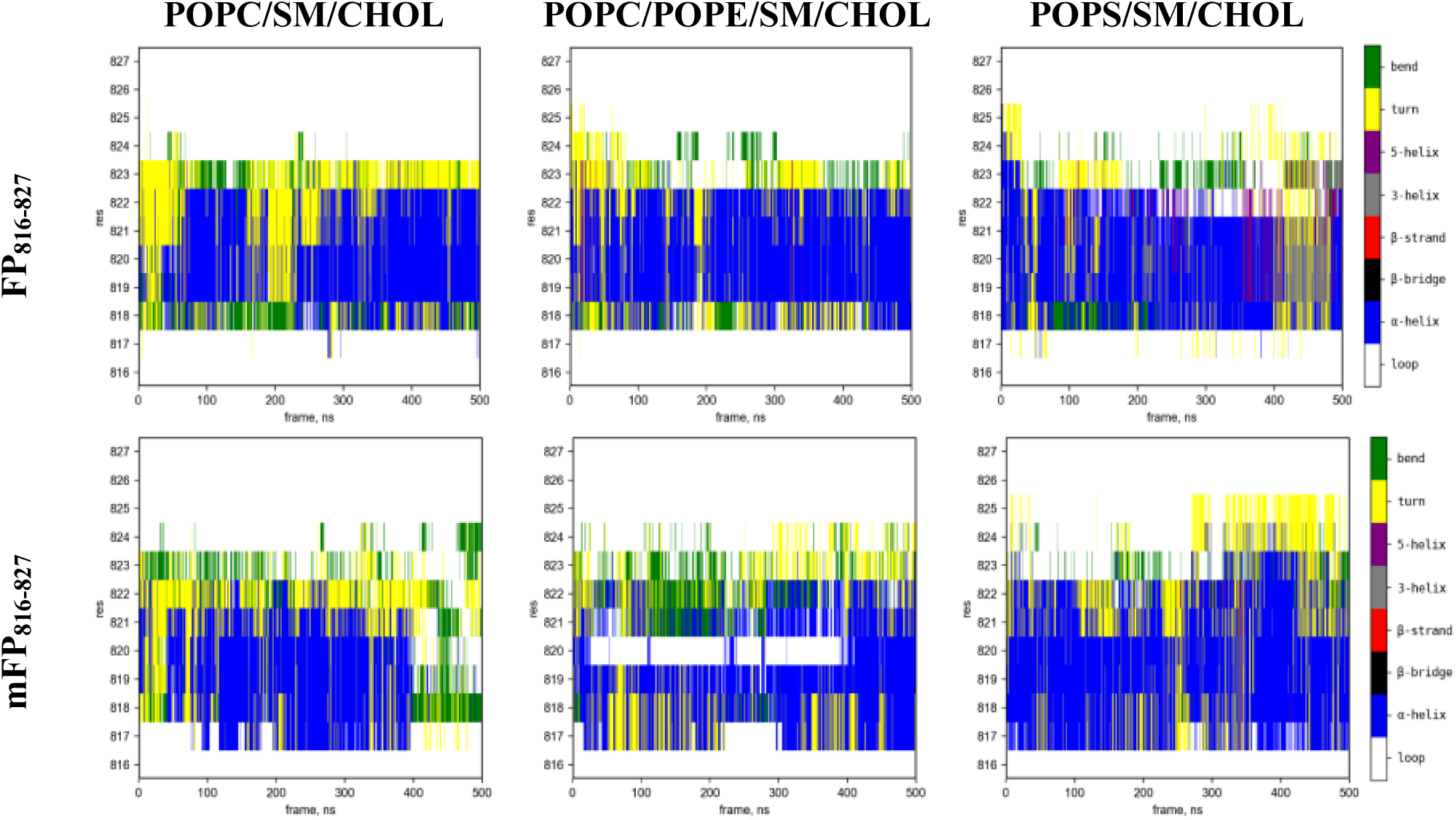
Secondary structure of FP_816-827_ (above) and mFP_816-827_ (below) averaged over all the replicas for each bilayer composition in which binding occurs (4 for POPC/SM/CHOL (60/20/20 mol.%) and POPC/POPE/SM/CHOL (30/30/20/20 mol.%), 3 for POPS/SM/CHOL (60/20/20 mol.%)) presented as heatmaps. Residue numbers are plotted against time frames of the trajectories, while the secondary structure is coloured in accordance with standard DSSP values (see colour bar on the right).

The results of simulations also revealed that in all lipid systems tested the peptides did end up partially embedded in the membrane. However, the extent to which FP_816-827_ and mFP_816-827_ disturb the lipid packing differs markedly. The values for displaced volume (i.e., occupied by the peptide wedged between lipid molecules) are distinctly higher for FP_816-827_ (Table 2). This is possibly due to the LLF motif not only being bulkier per se, but also carrying larger hydrophobic moieties capable of interacting with non-polar portions of the lipids. In agreement with these observations are the data on protein-lipid hydrophobic contacts (Table 2), indicating how strongly the peptides interact with the hydrophobic stratum of the bilayer. The mFP_816-827_ engages in such contacts much more poorly than FP_816-827_, aligning with its more superficial localization in the membrane.

**Table 2.** Estimated displaced volume (extvol) values and the number of protein-lipid hydrophobic (nphob) per frame in all-atom simulations.

| peptide | bilayer lipid composition | MEAN<br>0-500 ns |  | MEAN<br>0-250 ns |  | MEAN<br>250-500 ns |  |
| --- | --- | --- | --- | --- | --- | --- | --- |
|  |  | extvol, Å<sup>3</sup> | nphob | extvol, Å<sup>3</sup> | nphob | extvol, Å<sup>3</sup> | nphob |
| FP<sub>816-827</sub> | POPC/SM/CHOL | 125.4 | 120.4 | 130.7 | 124.4 | 120.1 | 116.5 |
|  | POPC/POPE/SM/CHOL | 125.1 | 121.8 | 130.2 | 124.2 | 120.0 | 119.3 |
|  | POPS/SM/CHOL | 126.5 | 131.3 | 122.3 | 127.9 | 130.7 | 134.6 |
| mFP<sub>816-827</sub> | POPC/SM/CHOL | 88.0 | 82.3 | 90.2 | 83.0 | 85.9 | 81.6 |
|  | POPC/POPE/SM/CHOL | 75.7 | 77.2 | 70.2 | 74.8 | 81.2 | 79.5 |
|  | POPS/SM/CHOL | 76.6 | 82.9 | 82.6 | 84.3 | 70.5 | 81.4 |

However, as regards more superficial strata of the membrane, the behavior of FP_816-827_ differs between model bilayers. Most notably, the peptide, via E819 and D820, and slightly via T827 engages in more significant non-covalent interactions (hydrogen bonds and salt bridges) with the POPE and POPS in POPC/POPE/SM/CHOL and POPS/SM/CHOL bilayers, respectively (Figure 4). This process does not take place in POPC/SM/CHOL system. For the mFP_816-827_ the picture is similar, with the exception of less pronounced binding of S816 to POPE compared to the case of FP_816-827_. Based on the study [35], which investigated the fusogenic activity of the FP1 domain of the S protein of SARS-CoV, characterized by high conservation and similarity to the homologous region in SARS-CoV-2, it can be hypothesized that residues I818 and K825 are also critically important for fusion. Substitution of these residues with alanine completely abolished cell-cell fusion and, in the case of K825, reduced the infectivity of pseudotyped virions. However, it is noteworthy that there are SARS-CoV-2 strains in which I818 is replaced by V [40]. This suggests that the volume of the amino acid residue radical within the fusion peptide also plays an important role in fusion. Overall, the FP of the S protein of SARS-CoV exhibits one of the lowest mutation densities [40].

**Figure 4.**
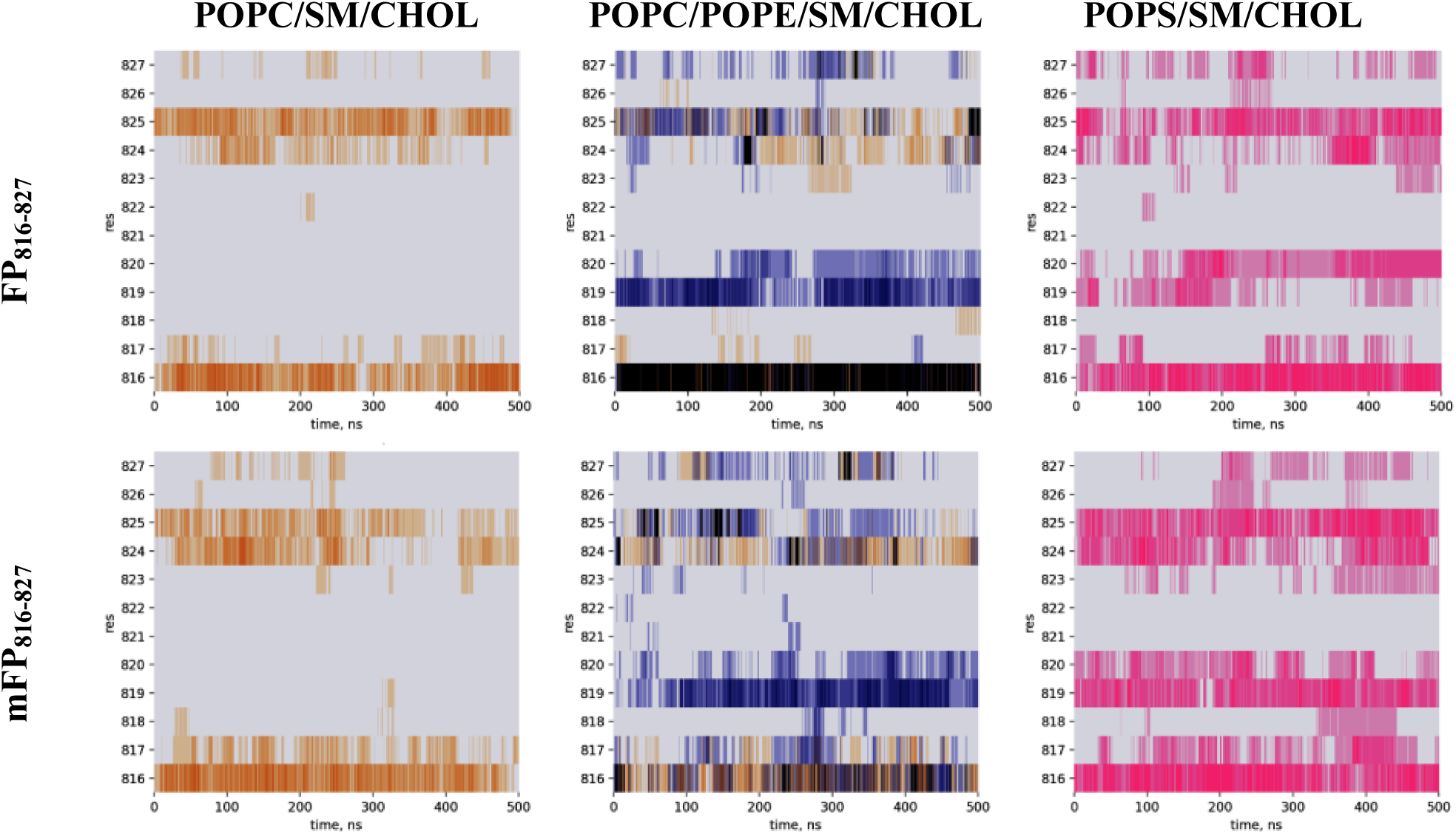
Hydrogen bonds engaged in by FP_816-827_ (above) and mFP_816-827_ (below) with model membranes. The heatmaps are superposed for all replicas run for each bilayer composition. Residue number is plotted against trajectory frame number, and the presence of hydrogen bonds with POPC, POPE and POPS is coloured yellow, purple and pink, respectively. The heatmap cell is coloured grey if no hydrogen bonds are detected.

We also analyzed how the binding of peptide fragments to lipid heads via hydrogen bonds affects the number of lipid molecules near the peptides, i.e., the formation of densely packed mini-domains in its own vicinity. This can be seen in graphs in which the distance between the peptide’s carboxyl groups and the cationic moieties in the lipids is plotted against the number of lipids located within such a radius from the peptide (Figure 5). Indeed, this number of lipids is greater at small distances from the peptide in POPC/POPE/SM/CHOL and POPS/SM/CHOL systems. At large distances, this trend persists only for POPC/POPE/SM/CHOL.

**Figure 5.**
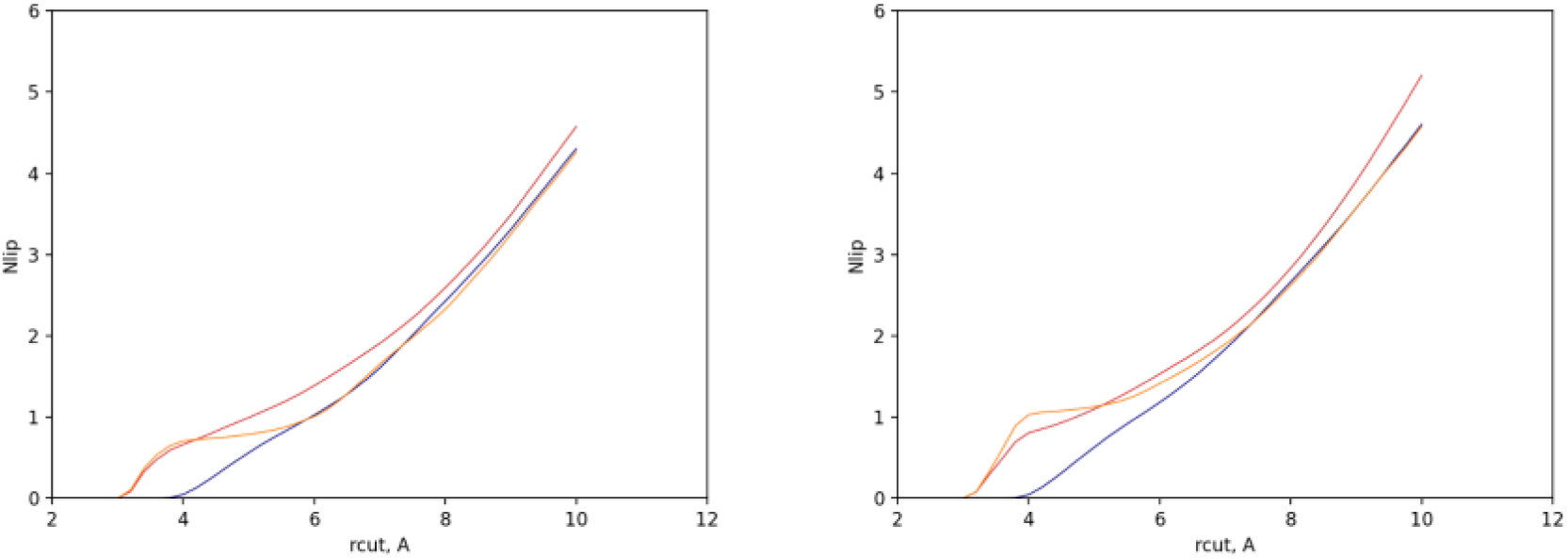
Number of lipids in the vicinity of FP_816-827_ (left) and mFP_816-827_ (right). The plots represent the number of lipids found within a given radius from the peptide. The curve for each bilayer composition has been averaged for all the replicas run. Blue, red, and orange curves are related to POPC/SM/CHOL (60/20/20 mol.%), POPC/POPE/SM/CHOL (30/30/20/20 mol.%), and POPS/SM/CHOL (60/20/20 mol.%) lipid bilayers respectively.

Important, that opposite processes unfold in different strata of the bilayer. While the lipid tails in the hydrophobic core are pushed apart, as the peptide penetrates this portion of the membrane, the lipid heads are, on the other hand, pulled together by the peptide. This could indicate that the bilayer is indeed disturbed, with abnormal patterns of lipid packing. It should be noted that the pushing apart of the lipid tails when the heads are pulled together under the influence of the FP_816-827_ is similar to an increase in negative spontaneous curvature. In the case of the mFP_816-827_ mutant this tendency would obviously be less pronounced, given its milder impact on the hydrophobic core (Table 2), and might determine its low fusogenic activity (Table 1). The more pronounced effect of the FP_816-827_ on the clustering of lipid heads at large distances (Figure 5) and on the number of hydrophobic interactions in POPC/POPE/SM/CHOL membranes (Table 2) might explain the increase in *IF* values induced by FP_816-827_ in POPE-enriched membranes compared to POPC/SM/CHOL system (Table 1). The number of hydrophobic interactions in POPS/SM/CHOL bilayers is even larger than in POPC/SM/CHOL and POPC/POPE/SM/CHOL membranes (Table 2), but unique nonmonotonic dependence of number of lipids located in FP_816-827_ vicinity on the distance from peptide might be related to decrease in ability of FP_816-827_ to fuse POPS/SM/CHOL liposomes compared to other tested lipid systems (Table 1). Considering all the obtained results, several factors contributing to the initiation of fusion can be proposed. When incorporated into the membrane, FP_816-827_ exerts a disordering effect (Table 3), which results in an increase in area per lipid and a decrease in the acyl chain order in the contacting monolayer. To compensate the resulting asymmetry, the opposite monolayer tends to be more ordered [41, 42]. The consequence of these perturbations is the emergence of localized protrusions in the upper monolayer characterized by a high positive spontaneous curvature. It is known that the formation of such protrusions reduces the contact area between membranes and thus lowers the hydration repulsion [43]. Another factor promoting fusion is the bending of acyl chains. Interestingly, our results are in agreement to molecular modeling studies, demonstrated the ability of fusion peptides from influenza and HIV to induce the repositioning of lipid tails and facilitate the intrusion of lipid heads [44,45]. Given the similarity of the results obtained with various fusion peptides, it is tempting to suggest that FP_816-827_ might exhibit similar activity. It is also noteworthy that all the aforementioned viral peptides belong to class I fusion proteins. This might indicate the existence of common fusion mechanisms characteristic of all proteins within the class. These mechanisms highlight the complex interplay between peptide conformation, lipid interactions, and membrane physical properties in driving the fusion process, providing valuable insights into the functionality of type I fusion proteins.

**Table 3.** The effects of FP_816-827_ and mFP_816-827_ on the thermotropic behavior of various phosphatidylcholines.

| peptide | ratio | DTPC (13:0) |  |  | DMPC (14:0) |  |  | DSPC (18:0) |  |  |
| --- | --- | --- | --- | --- | --- | --- | --- | --- | --- | --- |
| | | $\Delta T_m$, °C | $\Delta \Delta T_b$, °C | $\Delta \Delta H$, kcal/mol | $\Delta T_m$, °C | $\Delta \Delta T_b$, °C | $\Delta \Delta H$, kcal/mol | $\Delta T_m$, °C | $\Delta \Delta T_b$, °C | $\Delta \Delta H$, kcal/mol |
| FP<sub>816-827</sub> | 50:1 | -0.1 ± 0.1 | 0.8 ± 0.1 | -3.3 ± 0.4 | 0.1 ± 0.1 | 0 | -0.5 ± 0.2 | 0 | 0 | 0 |
|  | 25:1 | -0.2 ± 0.1 | 0.9 ± 0.2 | -4.4 ± 0.5 | 0.1 ± 0.1 | 0.1 ± 0.1 | -2.1 ± 0.4 | 0 | 0 | 0 |
|  | 10:1 | -0.4 ± 0.2 | 1.0 ± 0.3 | -7.9 ± 0.7 | 0.1 ± 0.1 | 0.2 ± 0.1 | -4.4 ± 0.6 | 0 | 0 | 0 |
| mFP<sub>816-827</sub> | 50:1 | 0 | 0 | 0 | 0.4 ± 0.1 | 0 | -0.1 ± 0.1 | 0 | 0 | 0 |
|  | 25:1 | 0.1 ± 0.1 | 0.2 ± 0.2 | -0.3 ± 0.1 | 0.6 ± 0.2 | 0 | -0.2 ± 0.1 | 0 | 0 | 0 |
|  | 10:1 | 0.3 ± 0.1 | 0.3 ± 0.1 | -0.4 ± 0.1 | 1.0 ± 0.2 | 0 | -0.3 ± 0.1 | 0 | 0 | 0 |

$\Delta T_p$ – the changes in the pre-transition temperature of the DPPC and DSPC in the presence of FP<sub>816-827</sub> and mFP<sub>816-827</sub>. The $T_p$ of pure DPPC and DSPC was equal to 34.8 ± 0.9 and 51.3 ± 0.4°C respectively. Pre-transition of DMPC was not analyzed.
$\Delta T_m$ , $\Delta \Delta T_b$ – the peptide-induced changes in the maximum temperature of the L<sub>β</sub>/L<sub>α</sub> transition of DTPC, DMPC, DPPC and DSPC and the temperature difference between the upper (onset) and lower (completion) boundary of the main phase transition of the lipids. The $T_m$ of unmodified DTPC, DMPC, DPPC and DSPC was equal to 13.9 ± 0.2 °C, 24.1 ± 0.2°C, 41.5 ± 0.1°C, and 54.9 ± 0.1°C respectively. $\Delta T_b$ of unmodified DTPC, DMPC, DPPC and DSPC was equal to 2.8 ± 0.3°C, 2.2 ± 0.5°C, 2.3 ± 0.3°C, and 2.0 ± 0.3°C respectively.
$\Delta \Delta H$ – the changes in the enthalpy of main transition of DTPC, DMPC, DPPC and DSPC in the presence of the peptides. The $\Delta H$ of unmodified DTPC, DMPC, DPPC and DSPC was equal to 20.0 ± 2.9, 15.8 ± 1.1, 16.7 ± 2.3 and 39.1 ± 3.4 kcal/mol respectively.

It should be especially noted that molecular dynamics approach had a limitation due to the fact that it did not include the simulation of processes of aggregation or oligomerization of FPs on the membrane surface, which is a characteristic feature of viral fusion peptides [46]. Moreover, choosing the box size in MD is debatable, as studies show that peptides folding depends on solvent volume in the simulation [47]. Many other parameters need to be noted while determining optimal box size, as little changes may lead to results discrepancy, when checking via other methods.

### Investigation of action of SARS-CoV-2 FP1 fragments on membranes of various composition using differential scanning microcalorimetry

Taking into account the influence of FP_816-827_ and mFP_816-827_ on lipid packing shown by *in silico* methods, differential scanning microcalorimetry was also performed.

Figure 6 demonstrates the thermograms of various phosphatidylcholines in the absence and presence of FP_816-827_ and mFP_816-827_ at different molar lipid:peptide ratios. Figure 7 shows the thermograms of various phosphatidylserines in the absence and presence of these peptides at different molar lipid:peptide ratios. The alignment of melting profiles of the subsequent heating scans attested to the high reproducibility of the results obtained (data not shown). The effects of peptides on the pre-transition of DPPC and DSPC (the transition from the gel (*L_β_*) to the ripple (*P_β’_*) phase) are shown on the insets Figure 6. To characterize the effects of FP_816-827_ and mFP_816-827_ on the thermotropic behavior of phosphatidylcholines we used following parameters: the alterations in the temperature of pre-transition (Δ*T_p_*), in the temperature of main transition to the liquid crystalline phase (*L_α_*) (Δ*T_m_*), in a width of the main peak (in a sharpness of main transition, ΔΔ*T_b_*)), and in the enthalpy of the main phase transition (in an area of the main peak, ΔΔ*H*). Δ*T_p_*, Δ*T_m_*, and ΔΔ*T_b_*, and ΔΔ*H* values of various lipids modified by FP_816-827_ and mFP_816-827_ are summarized in Table 2. Due to biphasic main transition of phosphatidylserines instead of Δ*T_m_* values the effects of the peptides were characterized by changes in the maximum temperature of first and second melting components (Δ*T_m1_*, Δ*T_m2_*). The parameters describing the influence of FP_816-827_ and mFP_816-827_ on thermotropic behavior of phosphatidylserines are shown in Table 4. Changes the heating-cooling hysteresis (in the difference between the temperatures of main phase transition at heating and cooling stage of DPPC in the presence of all tested peptides did not exceed 0.2°C.

**Figure 6.**
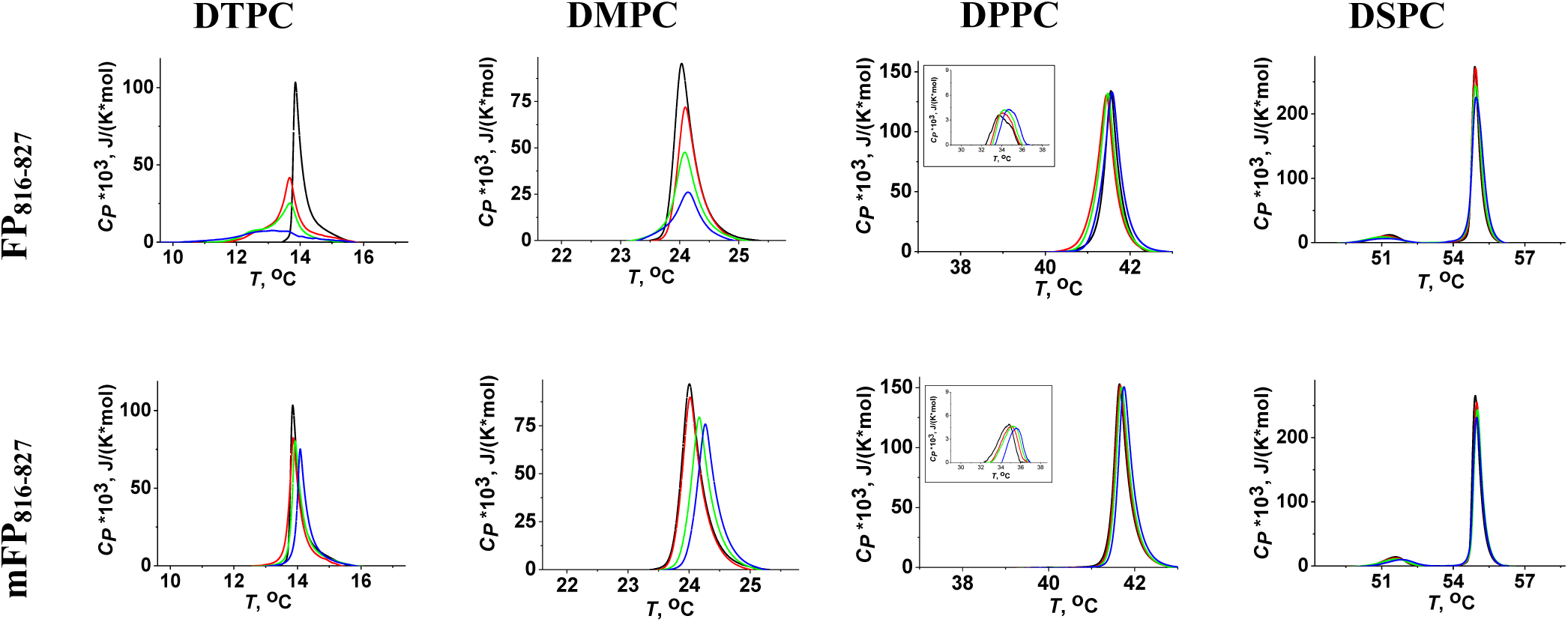
Heating thermograms of DTPC, DMPC, DPPC, and DSPC melting in the absence (control, *black curves*) and presence of FP_816-827_ and mFP_816-827 835_ at molar ratio of 50:1 (*red curves*), 25:1 (*olive curves*) and 5:1 (*blue curves*). *Insets:* Effects of peptides on the DPPC pre-transition.

**Figure 7.**
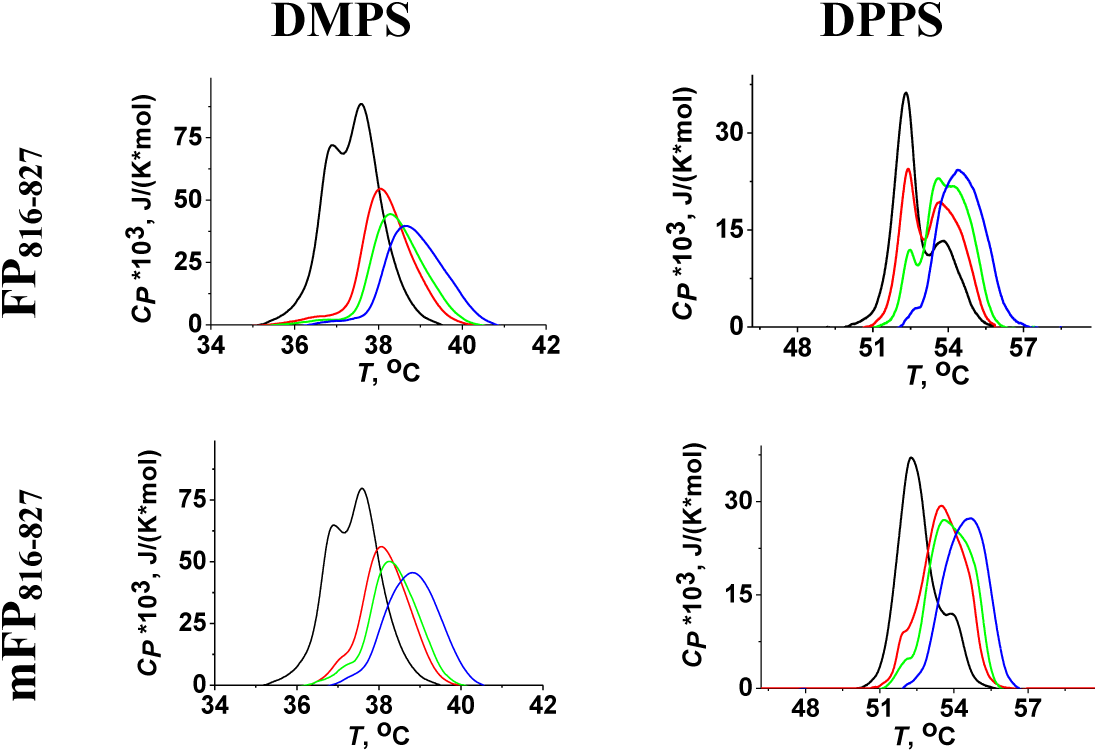
Heating thermograms of DMPS and DPPS melting in the absence (control, *black curves*) and presence of FP_816-827_ and mFP_816-827 835_ at molar ratio of 50:1 (*red curves*), 25:1 (*olive curves*) and 5:1 (*blue curves*).

**Table 4.** The effects of FP_816-827_ and mFP_816-827_ on the thermotropic behavior of various phosphatidylserines.

| peptide | ratio | DMPS (14:0) |  |  |  | DPPS (16:0) |  |  |  |
| --- | --- | --- | --- | --- | --- | --- | --- | --- | --- |
| | | $\Delta T_{m1}$, °C | $\Delta T_{m2}$, °C | $\Delta \Delta T_b$, °C | $\Delta \Delta H$, kcal/mol | $\Delta T_{m1}$, °C | $\Delta T_{m2}$, °C | $\Delta \Delta T_b$, °C | $\Delta \Delta H$, kcal/mol |
| FP<sub>816-827</sub> | 50:1 | $-0.2 \pm 0.1$ | $0.3 \pm 0.1$ | 0 | $-0.7 \pm 0.1$ | $0.1 \pm 0.1$ | 0 | $0.3 \pm 0.1$ | $-0.1 \pm 0.1$ |
| | 25:1 | $-0.1 \pm 0.1$ | $0.5 \pm 0.2$ | 0 | $-1.8 \pm 0.4$ | $0.2 \pm 0.1$ | $0.3 \pm 0.1$ | $0.4 \pm 0.2$ | $-0.2 \pm 0.2$ |
| | 10:1 | 0 | $0.8 \pm 0.1$ | 0 | $-1.9 \pm 0.3$ | —<sup>#</sup> | $0.8 \pm 0.2$ | $0.4 \pm 0.1$ | $-0.4 \pm 0.2$ |
| mFP<sub>816-827</sub> | 50:1 | $0.2 \pm 0.2$ | $0.3 \pm 0.1$ | 0 | $-2.1 \pm 0.3$ | $-0.1 \pm 0.1$ | $-0.2 \pm 0.1$ | $0.2 \pm 0.1$ | $-0.3 \pm 0.2$ |
| | 25:1 | $0.4 \pm 0.2$ | $0.5 \pm 0.1$ | 0 | $-2.2 \pm 0.2$ | $-0.2 \pm 0.1$ | 0 | $0.2 \pm 0.1$ | $-0.7 \pm 0.3$ |
| | 10:1 | —<sup>#</sup> | $0.7 \pm 0.2$ | 0 | $-2.5 \pm 0.2$ | —<sup>#</sup> | $0.6 \pm 0.2$ | $0.3 \pm 0.1$ | $-1.1 \pm 0.4$ |

$\Delta T_{m1}$ , $\Delta T_{m2}$ – the changes in the maximum temperature of first and second melting components of DMPS or DPPS in the presence of FP<sub>816-827</sub> and mFP<sub>816-827</sub>. The $T_{m1}$ and $T_{m2}$ of DMPS was equal to $36.9 \pm 0.1$ and $37.6 \pm 0.2$ respectively. The $T_{m1}$ and $T_{m2}$ of DPPS was equal to $52.3 \pm 0.1^\circ\text{C}$ and $53.8 \pm 0.1^\circ\text{C}$ respectively.
$\Delta \Delta T_b$ – the peptide-induced changes in the temperature difference between the upper (onset) and lower (completion) boundary of the main phase transition of DMPS and DPPS. The $\Delta T_b$ of unmodified DMPS and DPPS was equal to $3.1 \pm 0.5^\circ\text{C}$ and $5.8 \pm 0.4^\circ\text{C}$ respectively.
$\Delta \Delta H$ – the changes in the enthalpy of melting of DMPS and DPPS in the presence of the peptides. The $\Delta H$ of unmodified DMPS and DPPS was equal to $14.9 \pm 0.9$ and $12.6 \pm 2.5$ respectively.

Figure 6 and Table 3 clearly demonstrates that the effects of FP_816-827_ and mFP_816-827_ on the thermotropic behavior of phosphatidylcholines decreased as the length of the acyl tails increased. So, both fusogenic FP_816-827_ and inactive mFP_816-827_ demonstrated similar dose-dependent increase in *T_p_*of DPPC up to 0.6-0.8°C at the lipid:peptide molar ratio of 10:1 (Figure 6, *insets*, Table 3), while they were not able to modify the pre-transition of DSPC. The peptides had virtually no effect on the main phase transition of DPPC and DSPC (alteration in *T_m_* and Δ*T_b_* did not exceed 0.1°C, and ΔΔ*H* values were no more than 0.3°C (Figure 6, Table 3)). Wherein, unlike FP_816-827_, mFP_816-827_ was able to significantly affect melting of DMPC, a phosphatidylcholine with shorter (14:0) acyl chains than DPPC (16:0), by increasing *T_m_* on 1.0°C at the lipid:peptide molar ratio of 10:1 (Figure 6, Table 3). Wherein, FP_816-827_ decreased Δ*H* of DMPC (Figure 6, Table 3). Further decreasing in a phosphatidylcholine acyl chain length up to 13:0 (DTPC) was accompanied by significant decrease in *T_m_*on 0.4°C, increase Δ*T_b_* on 1.0°C, and pronounced reduction in Δ*H* on about 8 kcal/mol at the lipid:peptide molar ratio of 10:1 in the presence of FP_816-827._ Activity of mFP_816-827_ in DTPC compared to DMPC changes (Δ*T_m_* and ΔΔ*T_b_* are 0.3°C, and ΔΔ*H* values −0.4°C). At the same time, mFP_816-827_, unlike FP_816-827_, exerts a condensing effect (Figure 6, Table 3). The dependence of FP_816-827_ effects on the membrane hydrophobic thickness and FP_816-827_-induced decrease in DTPC melting temperature might be related to disordering effect of the peptide on acyl chains. These results are in agreement to data of molecular dynamics simulation showing the strong peptide interaction with the hydrophobic core of the POPC bilayer (Table 2).

Both peptides were more active against phosphatidylserines compared to phosphatidylcholines with tails of the same length (Figure 7, Table 4). Moreover, the effects of FP_816-827_ and mFP_816-827_ on the thermotropic behavior of phosphatidylserines increased as the length of the acyl tails increased in contrast to what was observed in the case of phosphatidylcholines. The interaction between the peptides and DPPS induced more substantial expansion of melting thermograms compared to phosphatidylcholines: FP_816-827_ and mFP_816-827_ established similar dose-dependent increase in Δ*T_b_* value of DPPC up to 0.3-0.4°C at the lipid:peptide molar ratio of 10:1 (Table 4). The peptides also decreased Δ*H* value of DPPS in a dose-dependent manner, and the efficiency of mFP_816-827_ was larger than of FP_816-827_ (–1.1 and –0.4 kcal/mol at lipid:peptide molar ratio of 10:1 respectively). Δ*H*-decreasing effects of the peptides enlarged as the length of the phosphatidylserine acyl tails decreased from 18:0 (DPPS) up to 16:0 (DMPS) (ΔΔ*H* values in the presence of FP_816-827_ and mFP_816-827_ were equal to –1.9 and –2.5 kcal/mol at lipid:peptide molar ratio of 10:1 respectively) (Table 4). To evaluate the peptide-induced changes in the transition temperature of distinct melting components, the peaks in the thermograms of phosphatidylserines were deconvoluted (Figure S1, Figure S2, Supplementary materials). Decomposition analysis revealed that independently on length of phosphatidylserine fatty acid chains and the peptide type the transition temperature of low-melting component (*T_m1_*) in the presence of the peptides changed slightly (Δ*T_m1_*values did not exceed 0.4°C), while the characteristic temperature of high-melting component (*T_m2_*) increased on 0.6-0.8°C at the lipid:peptide molar ratio of 10:1 (Table 4). Moreover, the component with a higher melting point became predominant compared to the control case, where the component with a lower melting point had a greater contribution (Figure 7, Figure S1, Figure S2, Supplementary materials). This effect was dose-dependent, and at the lipid:peptide molar ratio of 10:1 the component with a lower melting point was not observed at all. The phenomenon of biphasic melting of phosphatidylserines, the presence of low-melting and high-melting component, is associated with the difference in the protonation state of the carboxyl group [48]. Moreover, it was argued in [49] that low-melting and high-melting points correlate with degrees of lipid hydration. In other words, a higher melting point corresponds to a lower degree of lipid headgroup hydration. Thus, the predominance of a high component in the presence of the peptides (Figure S1, Figure S2, Supplementary materials) indicates that FP_816-827_ and mFP_816-827_ sorption on the membranes composed of phosphatidylserines leads to their dehydration, more probably, via electrostatic interactions. The dehydration of membrane surface might promote the membrane fusion [50–52].

It is known that, the functions of protein molecules are regulated not only by the polar part of lipids but also by their hydrocarbon segments. Bilayer thickness is one of the most crucial factors that can modulate the activity of various membrane proteins [53–56]. A mismatch between the length of the protein’s hydrophobic part and the hydrophobic thickness of the bilayer can affect the tilt angle of the protein when intercalating into the membrane, while the lipids in the membrane can also experience deformation, either stretching or compressing, to mitigate hydrophobic mismatch [57]. The length of the hydrophobic core of the membrane also influences the kinetics of insertion and the secondary structure of reconstituted peptides and proteins [58–61]. The differential scanning microcalorimetry data obtained with phosphocholines of varying tail lengths indicated that the effects of FP_816-827_ also depended on the thickness of the membrane’s hydrocarbon core. So, FP_816-827_ did not affect the thermotropic characteristics of the longer-chain lipid, DSPC (18:0), while it had a pronounced disordering effect on the shorter-chain lipid, DTPC (13:0) (Table 3). The hydrophobic thickness of DSPC and DTPC at 60°C is estimated as 31.9 and 22.4 Å [62]. The first way to explain the dependence of FP_816-827_ action on the lipid hydrocarbon chain is related to difference in peptide deformation to match the hydrophobic thickness of the bilayers of various composition. Peptide-induced disturbance of the lipid microenvironment might be altered by its deformation mode. Another way is associated to stronger van der Waals forces between lipid with longer chains [57], which increases the cost of peptide penetration.

## Materials and Methods

### Materials

Calcein, HCl, NaCl, CaCl_2,_ EDTA, HEPES, NaOH, sorbitol, Triton X-100, dimethylsulfoxide (DMSO), Sephadex G-50, were purchased from Sigma-Aldrich Company Ltd. (Gillingham, United Kingdom).

Lipids: 1-palmitoyl-2-oleoyl-*sn*-glycero-3-phosphocholine (POPC), 1,2-dioleoyl-*sn*-glycero-3-phosphocholine (DOPC), 1,2-ditridecanoyl-*sn*-glycero-3-phosphocholine (DTPC), 1,2-dimyristoyl-*sn*-glycero-3-phosphocholine (DMPC), 1,2-dipalmitoyl-*sn*-glycero-3-phosphocholine (DPPC), 1,2-distearoyl-*sn*-glycero-3-phosphocholine (DSPC), 1-palmitoyl-2-oleoyl-*sn*-glycero-3-phosphoethanolamine (POPE), 1-palmitoyl-2-oleoyl-*sn*-glycero-3-phospho-L-serine (POPS), 1,2-dimyristoyl-*sn*-glycero-3-phospho-L-serine (DMPS), 1,2-dipalmitoyl-*sn*-glycero-3-phospho-L-serine (DPPS), sphingomyelin (brain, porcine) (SM), cholesterol (CHOL), and 1,2-dipalmitoyl-sn-glycero-3-phosphoethanolamine-N-(lissamine rhodamine B sulfonyl) (ammonium salt) (Rh-DPPE) were obtained from Avanti Polar Lipids^®^ (Avanti Polar Lipids, Inc., USA).

Fusion peptide (FP) fragment of SARS-CoV-2 of 98% purity were synthesized by IQ Chemical (Saint-Petersburg, Russia). The peptides were synthesized by the standard solid phase method. The amino acid sequence of the samples was confirmed by MALDI-TOF mass spectrometry. Peptides were solubilized in DMSO at room temperature (25 ± 2°C). The choice of peptide was based on the work of Lai and Fried [63]. The peptide which is homologous to region 816-827 of the native SARS-CoV-2 FP with the highly conserved LLF motif responsible for membrane fusion [6,34] replaced for AAA [35], was used as a negative control. Namely, _816_SFIEDLLFNKVT_827_ (S2,FP1; FP_816-827_) and _816_SFIEDAAANKVT_827_ (S2,FP1; mFP_816-827_) sequences were used in the study.

### Membrane Fusion Assay

Large unilamellar vesicles were employed with a diameter of 100 nm, prepared through extrusion using an Avanti Polar Lipids® mini-extruder (Avanti Polar Lipids, Inc., USA). 3 mM of POPC/SM/CHOL (60/20/20 mol.%), POPC/POPE/SM/CHOL (30/30/20/20 mol.%), POPS/SM/CHOL (60/20/20 mol.%) mixtures in chloroform were placed in glass tubes and dried under a gentle stream of nitrogen for 5 minutes. The resulting dried lipid films were then resuspended in a 300 μl of buffer containing 35 mM calcein, 10 mM HEPES at pH 7.4. Subsequently, the suspension underwent five freeze/thaw cycles and was passed through a polycarbonate membrane 13 times. The removal of calcein located outside the vesicles was accomplished through gel filtration using Sephadex G-50, with the replacement buffer comprising 150 mM NaCl, 10 mM HEPES at pH 7.4. Liposomes were stored on ice, shielded from light, for a period of 2-3 weeks.

Given that calcein fluoresces weakly at millimolar concentrations, the intensity of its fluorescence in surrounding media can be used to assess the dye leakage during liposome fusion [31]. Calcein fluorescence was observed (excitation at 490 nm; emission at 520 nm) using a Fluorat-02-Panorama spectrofluorimeter (Lumex, St. Petersburg, Russia) equilibrated at 25 °C.

The fusion reaction was initiated by adding 150 μM of synthetic peptides, FP_816-827_ or mFP_816-827,_ to a suspension of vesicles with various lipid compositions.

The relative fluorescence of calcein (*IF, %*) representing the percentage of fused liposomes was calculated using the equation:

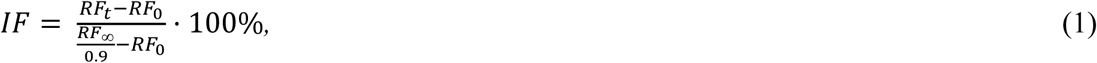

where *RF_0_*, *RF_t_* и *RF* are the fluorescence intensity of calcein at different times: 0, *t* and *t →* ∞, respectively. *RF* was taken to be the value after adding Triton X-100, which caused complete disengagement of calcein. To account the sample dilution, a factor of 0.9 was introduced.

*IF*-values were averaged from 2-5 independent experiments for all tested systems (mean value ± standard error) (*p* ≤ 0.05).

### Differential Scanning Microcalorimetry of Phosphatidylcholine and Phosphatidylserine

Giant unilamellar vesicles were obtained through electroformation using Vesicle Prep Pro® (Nanion Technologies, Munich, Germany) following the standard protocol (3 V, 10 Hz, 1 h) with heating to 25°C, 35°C, 55°C, and 65°C for DTPC, DMPC, DPPC (of DMPS), and DSPC (or DPPS), respectively. The suspensions contained 2.5 mM lipids and buffered with 5 mM HEPES at pH 7.4. The heating/cooling rate of the samples was 0.2 and 0.3°C/min, respectively. µDSC 7EVO microcalorimeter (Setaram, Caluire-et-Cuire, France) was used. Peptides (FP_816-827_ and mFP_816-827_), were added to the liposome suspension to achieve lipid:peptide ratios of 50:1, 25:1 and 10:1. Control samples remained unmodified. The results were analyzed using the Calisto software package (Setaram, Caluire-et-Cuire, France).

For the analysis of lipid melting thermograms, the following parameters were used:

1. The alteration in pre-transition temperature (changes in the temperature of the gel (*L_β_*) to ripple phase (*P_β_’*) transition, Δ*T_p_*). The effects of the tested peptides on the pretransition of DPPC and DSPC were analyzed.
2. The changes in the melting temperature (changes in the temperature of the main transition to a fluid phase (*L_α_*), Δ*T_m_*) and the width of the corresponding peak on the thermogram (*L_β_/L_α_* phase transition sharpness, ΔΔ*T_b_*)). Due to biphasic main transition of phosphatidylserines instead of Δ*T_m_*values the effects of the peptides were characterized by changes in the maximum temperature of first and second melting components (Δ*T_m1_*, Δ*T_m2_*).
3. The alteration in enthalpy of the main phase transition (an area of the main peak, ΔΔ*H*).
4. The changes in the hysteresis (in the temperature difference between the heating and cooling stages within one scan.

To confirm the reversibility of lipid thermal transitions, the difference in transition temperatures between the first and second heating repeated scans was utilized. The results were presented as the means of three independent experiments with several subsequent scans (mean value ± standard error) (p ≤ 0.05).

### Molecular dynamic simulations

The FP_816-827_ model was based on the said fragment from the pre-fusion SARS-CoV-2 spike trimer (PDB ID 6XR8) [10], while the mutations to create the mFP_816-827_ model were introduced using the Modeller9.19 package [64]. In order to identify states in which the peptide binds to the membrane, both models were converted to the coarse-grained representation using the martinise.py script [65,66]. Four replicas were made for each peptide in each of the following model bilayers: POPC/SM/Chol, POPC/POPE/SM/Chol, POPS/SM/Chol. In each replica the peptide was arbitrarily placed in the aqueous environment at a distance from the membrane using the *insert-molecules* utility in the GROMACS package [67]. MD simulations were performed using GROMACS 2021.5. Steepest descent energy minimization was performed for 2000 to 5000 steps followed by 1000ns MD simulation in the Martini force field [65,66] with an integration step of 20 fs under 3D periodic boundary conditions. Na+ and Cl− ion parameters for counter ions were implemented. Simulations were performed at 330 K temperature and 1 bar pressure maintained using the V-rescale [68] and the Berendsen [69] algorithms with 1.0 and 12.0 ps relaxation parameters, respectively, and a compressibility of 3 × 10^−4^ bar^−1^ for the barostat. The protein, membrane lipids and solvent molecules were coupled separately. Bonds with an H atom were constrained via implementing LINCS [70].

CG simulations completed, systems where the peptide bound to the membrane were converted to all-atom representation to study the peptide/membrane interaction in greater detail. The peptide was converted to all-atom representation via an algorithm based on backward.py [71]. Refinement of the peptide conformation based on the structure obtained at the end of the CG simulations was made using additional so-called ExpRst constraints imposed on the distances between CA atoms (i) and on the distances (0.2 nm) between backbone H and O atoms, for which hydrogen bonds (force constant 500 kJ/nm^2^) had been detected (ii). All-atom simulations were carried out using the Amber14 force field [72] for the peptide and the Slipids model [73] for the lipids. All-atom MD simulations were performed using the GROMACS 2021.5 package [67]. An integration time step of 2 fs was used, and 3D periodic boundary conditions were imposed. The spherical cut-off function (12 Å) was used to truncate van der Waals interactions. Electrostatic interactions were treated using the particle mesh Ewald (PME) method [74] (real space cutoff 12 and 1.2 Å grid with fourth-order spline interpolation). The TIP3P water model was used [75], and Na+ and Cl− ion parameters for counter ions were implemented. Simulations were performed at 310 K temperature and 1 bar pressure maintained using the V-rescale [68] and the Parrinello–Rahman [76] algorithms with 0.5 and 10.0 ps relaxation parameters, respectively, and a compressibility of 4.5 × 10^−5^ bar^−1^ for the barostat. The protein, membrane lipids, and solvent molecules were coupled separately. Semi-isotropic pressure coupling in the bilayer plane and along the membrane normal was used in the simulations. Before the simulation runs, all systems were equilibrated over three steps with imposed ExpRst: (i) steepest descent energy minimization over 1000 steps; (ii) 100-ps MD NVT for relaxation of steric clashes resulting from resolution transformation with an integration step of 1 fs; (iii) 1-ns MD semi-isotropic NPT for the relaxation of the lipid bilayer with an integration step of 1 fs. Production runs were simulated for 0.5 μs. Bonds with an H atom were constrained via implementing LINCS [70].

Trajectories were analysed using in-house software.

## Acknowledgements

This research was supported by the Russian Foundation of Science (project # 22-15-00417).

## Competing interests

The authors declare no competing interests.

## Additional information

**Supplementary information.** The online version contains supplementary material available.

## Supplementary materials

**Table S1.** Mean RMSD values in all-atom simulations.

| peptide | bilayer lipid composition | MEAN RMSD | MEAN RMSD | MEAN RMSD |
| --- | --- | --- | --- | --- |
|  |  | 0-500 ns, nm | 0-250 ns, nm | 250-500 ns, nm |
| **FP<sub>816-827</sub>** | POPC/SM/CHOL | 0.27 | 0.25 | 0.28 |
|  | POPC/POPE/SM/CHOL | 0.27 | 0.25 | 0.29 |
|  | POPS/SM/CHOL | 0.28 | 0.27 | 0.29 |
| **mFP<sub>816-827</sub>** | POPC/SM/CHOL | 0.29 | 0.28 | 0.31 |
|  | POPC/POPE/SM/CHOL | 0.32 | 0.30 | 0.33 |
|  | POPS/SM/CHOL | 0.25 | 0.26 | 0.24 |

**Figure S1.**
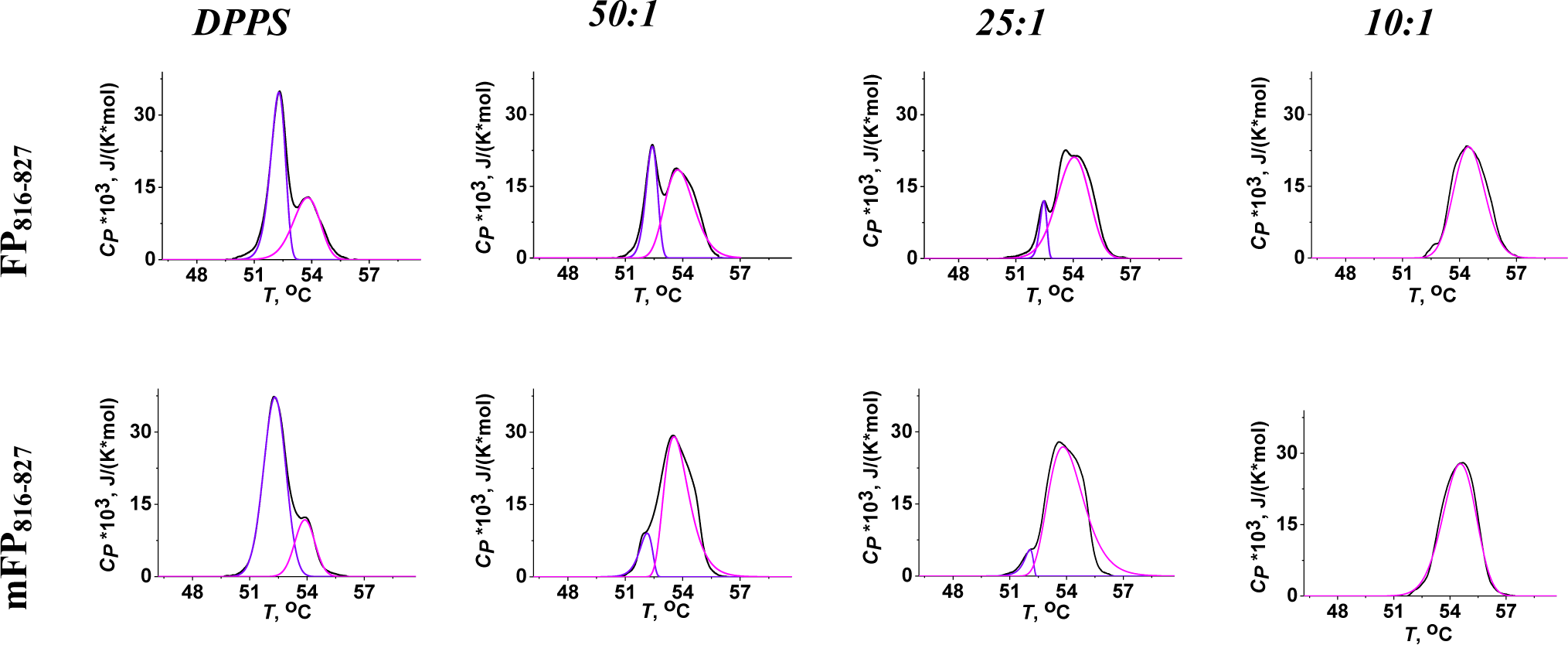
Deconvolution analysis of the main transition peak of DPPS in the presence of FP_816-827_ and mFP_816-827_ at different lipid:peptide molar of 50:1, 25:1 and 10:1. The parameters characterizing the peptide-induced changes in the individual components are summarized in Table 4.

**Figure S2.**
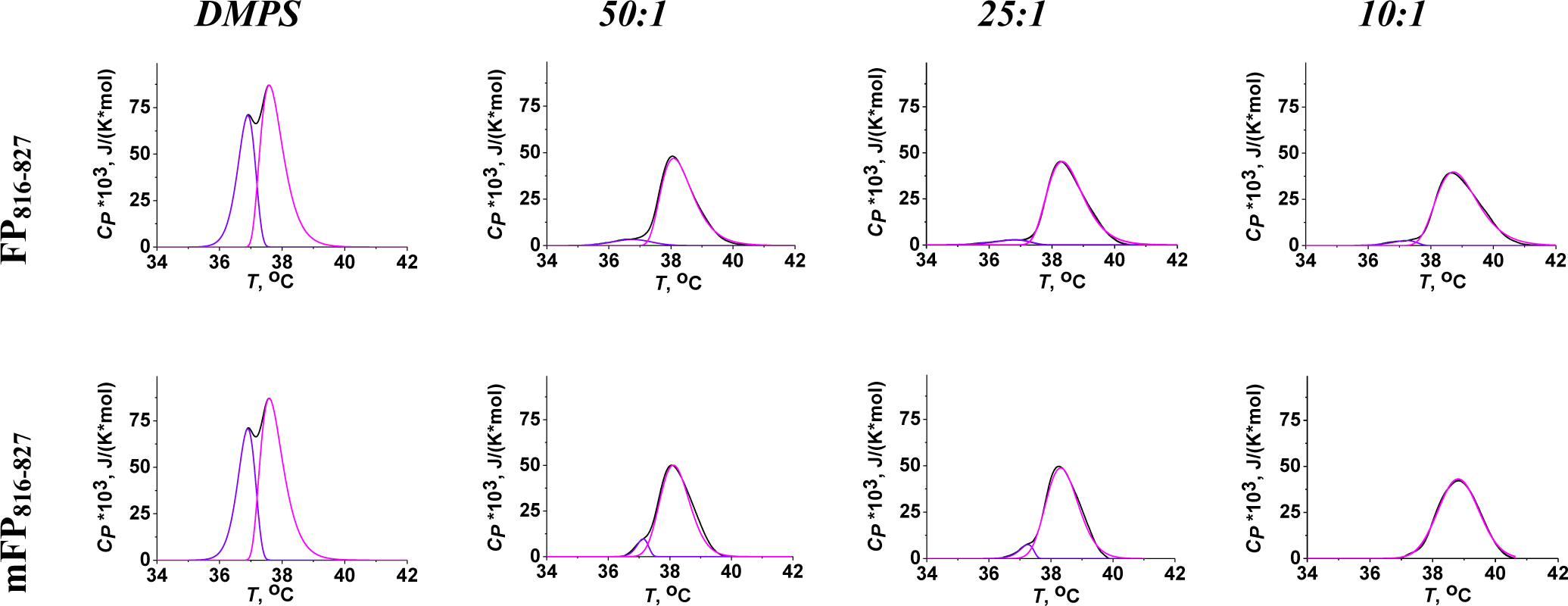
Deconvolution analysis of the main transition peak of DMPS in the presence of FP_816-827_ and mFP_816-827_ at different lipid:peptide molar of 50:1, 25:1 and 10:1. The parameters characterizing the peptide-induced changes in the individual components are summarized in Table 4.

## Notes

### Competing Interest Statement

The authors have declared no competing interest.

